# Growth-factor ligand functionalization enhances cellular and *in vivo* uptake of DNA nanodevices

**DOI:** 10.1101/2025.08.01.668157

**Authors:** Abhijit Biswas, Naveena A Hema, Krupa Kansara, Bhagyesh Parmar, Vivek Shekhar, Ashutosh Kumar, Sharad Gupta, Dhiraj Bhatia

## Abstract

Epidermal Growth Factor (EGF) and Transforming Growth Factor beta (TGF-β) are two important classes of growth factor that regulates cell growth, cell proliferation, differentiation, immune responses, and extracellular matrix generation. Using a short peptide-based ligand coupled to DNA nanocages, we present the enhanced internalization of a receptor-mediated peptide-DNA nanocage. We used here tdDNA as a delivery vehicle. Our study in cellular and in vivo showed excellent internalization, cell growth, and inhibition. We expect that a tdDNA-modified receptor-binding peptides could become a valuable scaffold for use as a cellular programming and regenerative material.

## 1. Introduction

The cellular environment is varied and constantly changing, offering both structural and biochemical assistance. Specifically, the extracellular matrix (ECM) influences the dynamics, spatial arrangement, and combined effects of signals to guide cellular behaviour^1–3^. Synthetic versions of this matrix for tissue repair must replicate these three traits within a unified system. In biological systems, components of ECM such as proteins, lipids, and DNA regulate the self-assembly, structural characteristics, and nanoscale display of bioactive signals^4–6^. Mimicking these traits in synthetic materials is essential for their integration with biological systems, for purposes that encompass therapeutic cargo delivery, target sensing and imaging, tissue engineering, and foundational research into biological mechanisms and forces^7^. Recently, two molecular platforms gaining popularity for the creation of self-assembling materials are synthetic peptides and DNA^8^. DNA, which adheres to the reliable Watson−Crick base pairing principles, offers nearly endless programmability for creating intricate nanostructures^1–13^, selectively immobilizing various ligands based on their sequences, and enabling reversible connections between two elements via strand-displacement reactions^9,4,11,12,14–2010^. Peptides, due to their extensive library of building blocks, exhibit a significant degree of variation and self-assembly characteristics, resulting in greater diversity and structural fluidity^11,12^. Consequently, DNA-peptide conjugates harness the advantageous properties of both components while also mitigating their individual limitations^3,7^. For instance, peptides can gain from the precise control of self-assembly offered by DNA nanotechnology, while DNA can be improved by the enhanced biological and mechanical stability provided by peptides. This makes peptide-DNA conjugates excellent candidates for investigation in medicine and therapy, as well as for the nanoscale manipulation and organization of bioactive substances.

A growth factor is a naturally occurring protein that functions as a signalling substance, promoting cell growth, healing of wounds, and, in some cases, differentiation of cells. Epidermal Growth Factor (EGF) mainly promotes the division of cells, whereas Transforming Growth Factor-beta (TGF-beta) has the potential to either suppress or promote cell growth based on the type of cell and the situation. EGF serves as a mitogen, fostering cellular growth, while TGF-beta can function as a tumor inhibitor at specific stages and as a tumor enhancer at other times. Balance between these two growth factor in body is essential for proper functioning^13–16,16,17^. In this study, we have designed a human EGF and TGF-β receptor-promoting peptide conjugated with tetrahedron DNA (td-DNA), a DNA origami for the successful internalization of receptor-promoting peptides inside the cells. Here we report the successful internalization of our engineered systems in vitro and in vivo model study.

## 2. Materials and methods

### 2.1 Materials and reagents

All amino acids and reagents for peptide synthesis were obtained from BLD Pharma, and Sigma Aldrich. The oligonucleotide strands (M1, M2, M3, M4, M4-Cy3) were sourced from Sigma. Dulbecco’s Modified Eagle Medium (DMEM), Fetal bovine serum (FBS), Penicillin-Streptomycin (PenStrep), and Trypsin-EDTA (0.25%), were acquired from Gibco. Nuclease-free water, ammonium persulfate, ethidium bromide, TEMED, Triton-X, paraformaldehyde, and adherent cell culture dishes came from Himedia. Tris-Acetate EDTA (TAE) and 40% Acrylamide/bis(acrylamide) solution were procured from GeNei. Magnesium chloride was ordered from SRL, India. SPDP (N-succinimidyl 3-(2-pyridyldithio) propionate) was bought from Invitrogen.

### 2.2. Synthesis of peptide

Peptides were synthesized using an in-house facility using the Fmoc-based solid-phase peptide synthesis strategy. The peptide was created on Fmoc-protected rink amide resin with a loading capacity of 0.8 mmol/g. The reaction was conducted on a scale of 100 mg. The resin was measured and allowed to swell in anhydrous DCM for 3 hours, then swelled in DMF for 3 hour. After draining, the resin was alternatively washed with DCM and DMF. The resin underwent Fmoc removal and the coupling of the initial amino acid through the subsequent procedures:

#### A. Fmoc Removal

The Fmoc was removed using 5% piperazine and DBU in DMF. This removal process was performed over three cycles with incubation durations of 12, 10, and 5 minutes, respectively. Following each removal, washes with DCM-DMF were conducted.

#### B. Amino Acid Coupling

The coupling of the Fmoc-protected amino acids was executed using the DIC/oxyma approach. The amino acid and oxyma were utilized in three equivalents, dissolved in DMF, and activated with three equivalents of DIC for 5 minutes while being gently shaken. Double coupling was performed to attach the activated amino acid to the growing peptidyl resin. Each coupling took place over the course of 1 hour. After the coupling process, the peptidyl resin underwent DCM-DMF washes. The procedures in Steps 1 and 2 were reiterated until the target peptide sequence was achieved. After synthesis, the peptidyl resin was rinsed with methanol. The peptide was detached from the resin using a cleavage cocktail made up of 92.5% TFA, 2.5% DODT, 2.5% TIPS, and 2.5% water for a duration of 3 hours with continuous vortex mixing. The resin was eliminated through filtration, and the resulting filtrate was concentrated. Peptides were precipitated under a nitrogen flow, reducing the total volume to approximately one-fifth. Following this, the peptides were treated with ice-cold diethyl ether and centrifuged at 8000 rpm for 30 minutes at a temperature of 4°C. The liquid supernatant was removed, and the resulting pellet was re-dissolved in a solution of 50% acetonitrile in water before being frozen and lyophilized. The resulting lyophilized powder was then stored at -20°C for future use. The characterization of the peptides was performed using LC-MS^18^.

### 2.3. Coupling of peptide and DNA oligo

The peptide modified with cysteine was coupled with the 5’ amino-modified M1 oligonucleotide using SPDP cross-linking. In summary, the amino-modified M1 strand at a concentration of 20 μM was incubated with 2 mM of SPDP in PBS-EDTA while continuously mixing for 1 hour at room temperature. A 7K Zeba spin desalting column was used to remove unreacted SPDP. The peptide was prepared in PBS-EDTA. Subsequently, M1-SPDP was incubated with 5 equivalents of the peptide in an equimolar amount to achieve a final working concentration of 10μM of M1 linked to the peptide. Unreacted peptide was eliminated through desalting.

### 2.4. Synthesis of DNA tetrahedron coupled with peptide

Four oligonucleotide strands (M1, M2, M3, M4) were utilized to create a DNA tetrahedron **(Table S1)**. The oligonucleotide strands were mixed with nuclease-free water to achieve a concentration of 100μM and heated to 70°C for 1 hour and 30 minutes. The M1 strand was coupled with the peptide. The remaining three strands were diluted to a working concentration of 10μM. The DNA tetrahedron (td) was assembled via a one-pot synthesis method, where all four oligonucleotide strands (M1-pep, M2, M3, M4) were added in equimolar amounts along with 2 mM MgCl2. This reaction took place at 95°C for 10 minutes and was immediately cooled to 4°C. The final concentration of td was set at 2.5 μM and stored at 4°C until it is needed for subsequent applications.

### 2.5. Electrophoretic Mobility Shift Assay

The Electrophoretic Mobility Shift Assay (EMSA) was employed to evaluate the formation of higher-order structures. A 10% Native-PAGE was utilized to conduct the EMSA. 5μl of the DNA td-pep samples were combined with 3 μl of loading buffer and 1.5μl of 6X loading dye. The Native gel was operated at a constant voltage of 70V for a duration of 120 minutes. The gel was then stained with 0.32 μg/ml of ethidium bromide for 10 minutes and visualized using the Gel documentation system (Biorad ChemiDoc MP Imaging System).

### 2.6. Cell culture

SUM-159 and RPE1 cells were grown in Dulbecco’s Modified Eagle Medium (DMEM) that was supplemented with 10% fetal bovine serum and 1% penicillin-streptomycin. The cells were incubated at 37°C in a humidified environment with 5% CO2. Typically, the cells were subcultured once they reached 85-95% confluency, and the media was replaced as necessary.

### 2.7. Cellular uptake studies

In the td-peptide uptake experiment, cells were plated at a density of 0.1 × 10^6^ cells per well on 12 mm coverslips and allowed to incubate for 24 hours. After washing the cells with 1X PBS, they were treated with either 100 nM of td or td-pep for 15 min in serum-free media at 37°C. Subsequently, the cells were washed three times with 1X PBS to eliminate any unbound or surface-attached td and td-pep nanostructures. The cells were then fixed using 4% paraformaldehyde at 37°C for 15 minutes. After that, the cells underwent three additional washes with 1X PBS, and the coverslips containing the cells were mounted onto glass slides using mowiol with DAPI to stain the nuclei.

### 2.8. Zebrafish Maintenance

Zebrafish (Danio rerio) (Assam wild-type) were raised from embryos to adults under controlled lab conditions, adhering to ZFIN guidelines. A light cycle of 14 hours on and 10 hours off was maintained at temperatures ranging from 26 to 28 °C. The fish were kept in aerated 20 L tanks with water parameters adjusted to replicate their natural habitat: a pH of 7 to 7.4, conductivity between 250 and 350 μS, tDS 220–320 mg L^−1^, salinity from 210 to 310 mg L^−1^, and dissolved oxygen levels exceeding 6 mg L^−1^, which were monitored using a PCD 650 multi-parameter device. Their diet included live Artemia fed twice daily and Aquafin basic flakes provided once a day. For breeding purposes, a ratio of 3 females to 2 males was employed, with embryos collected in E3 medium and incubated at 28.0 °C.

### 2.9. Confocal microscopy imaging

All confocal imaging of fixed cells was performed using the Leica TCS SP8 confocal laser scanning microscope. Images were obtained with a 63X oil immersion objective. The pinhole was maintained consistently at 1 Airy unit. The bit depth, laser power, and detector gain were held constant across various experimental conditions within a specific study. All images were captured at a resolution of 512 × 512. DAPI excited with a Diode 405 laser at 405nm, while Cy3 was excited at 561 nm with a DPSS laser, and Cy5 was excited at 633 nm with a He-Ne laser. The emission collection window for the detector was configured according to the emission spectra of the respective fluorophores. Z-stack imaging was conducted, and the number of z-steps was set to optimize the system. For imaging multiple fluorophores, sequential scanning was used.

### 2.10. Image Processing and Statistical Analysis

The analysis of the images was carried out using Fiji ImageJ software. Z-stacks were merged into 2D images using the z-projection method, specifically the maximum intensity projection approach. Background correction involved subtracting the average intensity of the background. Following this adjustment, both integrated density and raw integrated density were assessed. To conduct statistical comparisons, an unpaired, one-way ANOVA was used for two experimental groups, while analysis of variance (ANOVA) with Dunnett’s test was utilized for comparisons that included more than two experimental groups. All tests were performed under the assumption of normal distribution.

## 3. Results and Discussions

### 3.1 Engineering of peptide-DNA nanocages

The Epidermal Growth Factor (EGF) receptor (EGFR) and the Transforming Growth Factor-beta (TGF-beta) receptor are essential in regulating cell growth, proliferation, differentiation, and play a significant role in the onset of various diseases, including cancer. EGF stimulates cell proliferation, whereas TGF-beta serves as a strong inhibitor of cell growth and can also facilitate cell migration. Changes in expression or abnormal activation of these receptors can lead to the development and advancement of cancer. It is important to design hEGF and TGF-β promoting short peptides and its successful delivery in the affected cells. In this context we have chosen two short peptides (**Table 1, SI Figure 1-3**) which are the active site for promoting EGF and TGF-β growth factors in the body. To check their function we need to internalize them into cells. Currently lots of vehicles are used for delivery of short peptides in cells including polymers^19,20^, lipids^21,22^, cell-penetrating peptide (CPPs)^23–27^, DNA nanocages^7,28–32^ etc. The Bhatia group has conducted significant research on DNA origamis and found that tetrahedron DNA (tdDNA) holds tremendous promise for delivering cargo molecules into cells^7,6,21–39,28–32^. For conjugation of our chosen peptides with tdDNA, we have used amine functionalized tdDNA and Cys containing (-N terminus) peptides for SPDP crosslinking following the well-established protocol^33^. The successful conjugation of peptides with tdDNA has been confirmed by EMSA (Electrophoretic Mobility Shift Assay), with 10 % native gel The nanostructures of peptide-tdDNA conjugates obtained from AFM study with average diameter for AB1-tdDNA and AB2-tdDNA are 72.03 ± 22.8 nm and 79.63 ± 17.66 nm respectively (**Figure 1**).

**Table 1.** Designed peptide’s sequences.

| Peptide Code | Peptide Sequence |
| --- | --- |
| AB1 (TGF-β promoting)<sup>34</sup> | CGGGWSLD |
| AB2 (hEGF promoting)<sup>35</sup> | CGGGSCIPGESSDNCTALVQTEDNPRVAQ |

**Figure 1.**
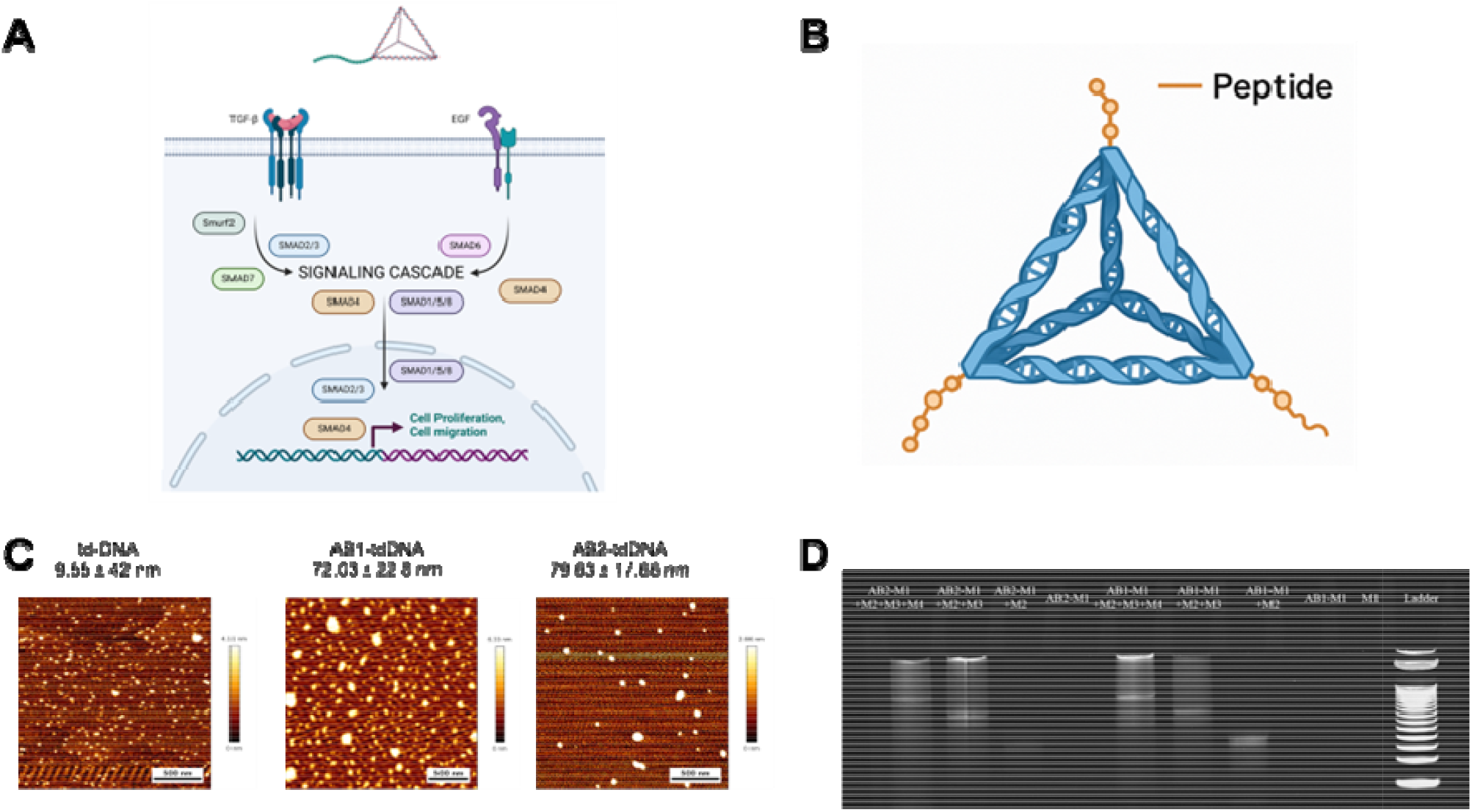
(A) Schematic of functional pathways of peptide-tdDNA conjugates in EGF and TGF-β promoting, (B), respresntative schematic of peptide conjugated tdDNA, (C) the morphology of td DNA and peptide-DNA conjugates obtained from AFM, and (D) gel electrophoresis showed the successful conjugation of the peptide with tdDNA.

### 3.2. Cellular internalization of peptide-tdDNA complexes

The cellular uptake study of the peptide-tdDNA was studied at 100 nM concentration on SUM-159 cells (a triple-negative breast cancer cell line) via confocal microscopy (**Figure 2**). We have incubated the peptide-tdDNA complex (Cy3 dye conjugated) for 15 min at 37°C in 5% CO_2_ incubator prior to fixing and confocal microscopy analysis. We have observed a high-intensity red signal inside the cells, and it is also confirmed by its quantification study using ImageJ software. These results suggest that our designed system successfully internalizes the receptor-promoting peptides in the cells.

**Figure 2.**
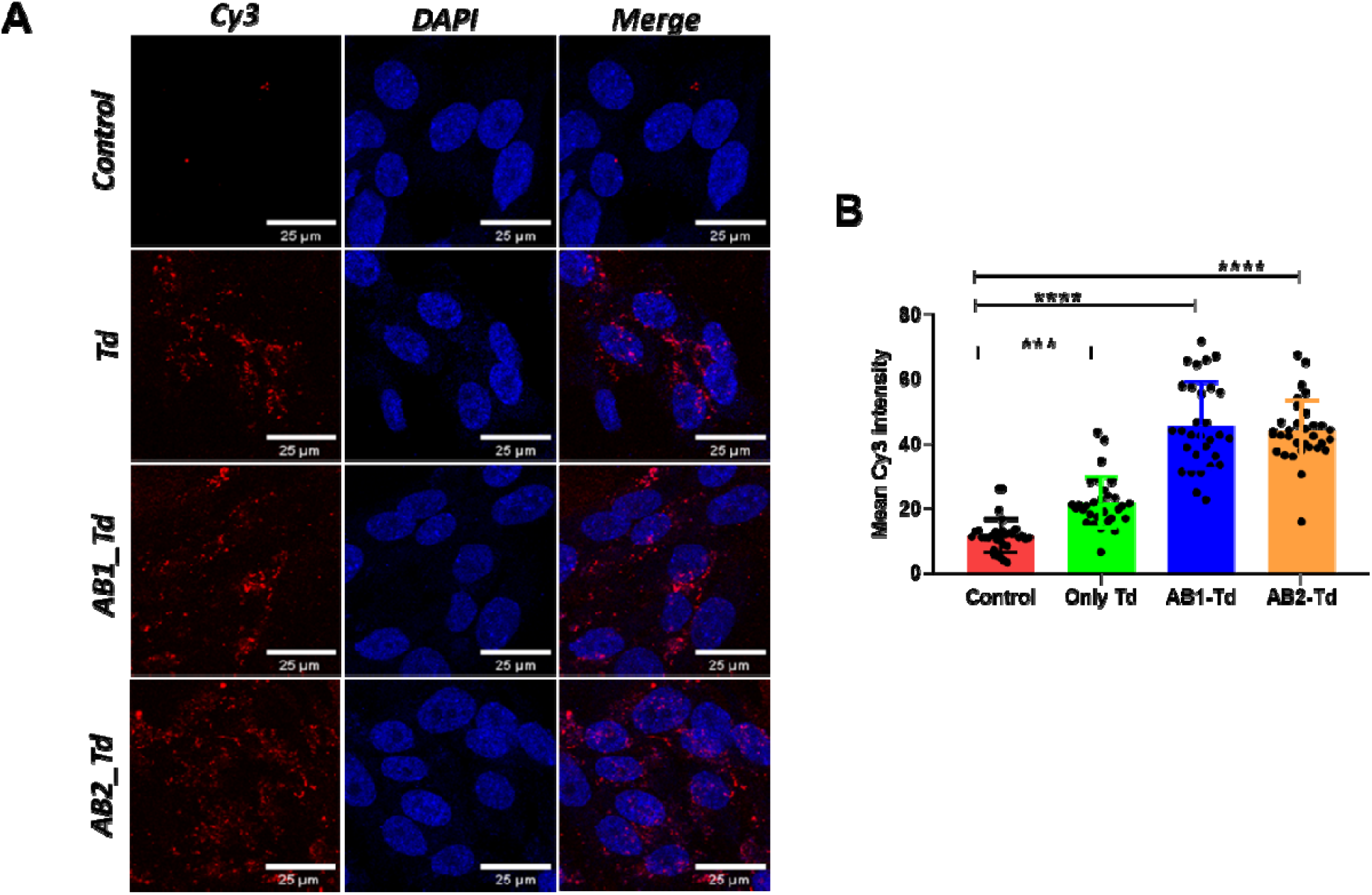
Intracellular distribution of peptide-tdDNA-Cy3 complex at 100 nM final concentration, after 15 min of incubation in SUM-159 cell line. Nuclei are stained with DAPI (blue), red represents the Cy3 labelled tdDNA **(A)**, the scale bar represent 50 µM. In the panel (**B)** it shows the quantification of internalization of Cy3 conjugated tdDNA in SUM-159 cells. * p-value < 0.05, ** p-value < 0.005, *** p-value < 0.0005, **** p-value < 0.0001.

### 3.3. Wound healing assay

We have applied our peptide-tdDNA complexes (100 nM concentration) to RPE1 cells (Retinal Pigment Epithelial-1 cells). Before applying the complex, we have to make a scratch on the cells at its confluency. The growth of cells was checked via an optical microscope at scheduled time points (0 h, 3h, 6h, 24h, and 30h). A significant observation was made from this study that cells which was treated with EGF-promoting peptide, proliferate very fastly compared to that of cells with TGF-promoting peptide, which also aligned to the other previous studies^36–41^ (**Figure 3**).

**Figure 3.**
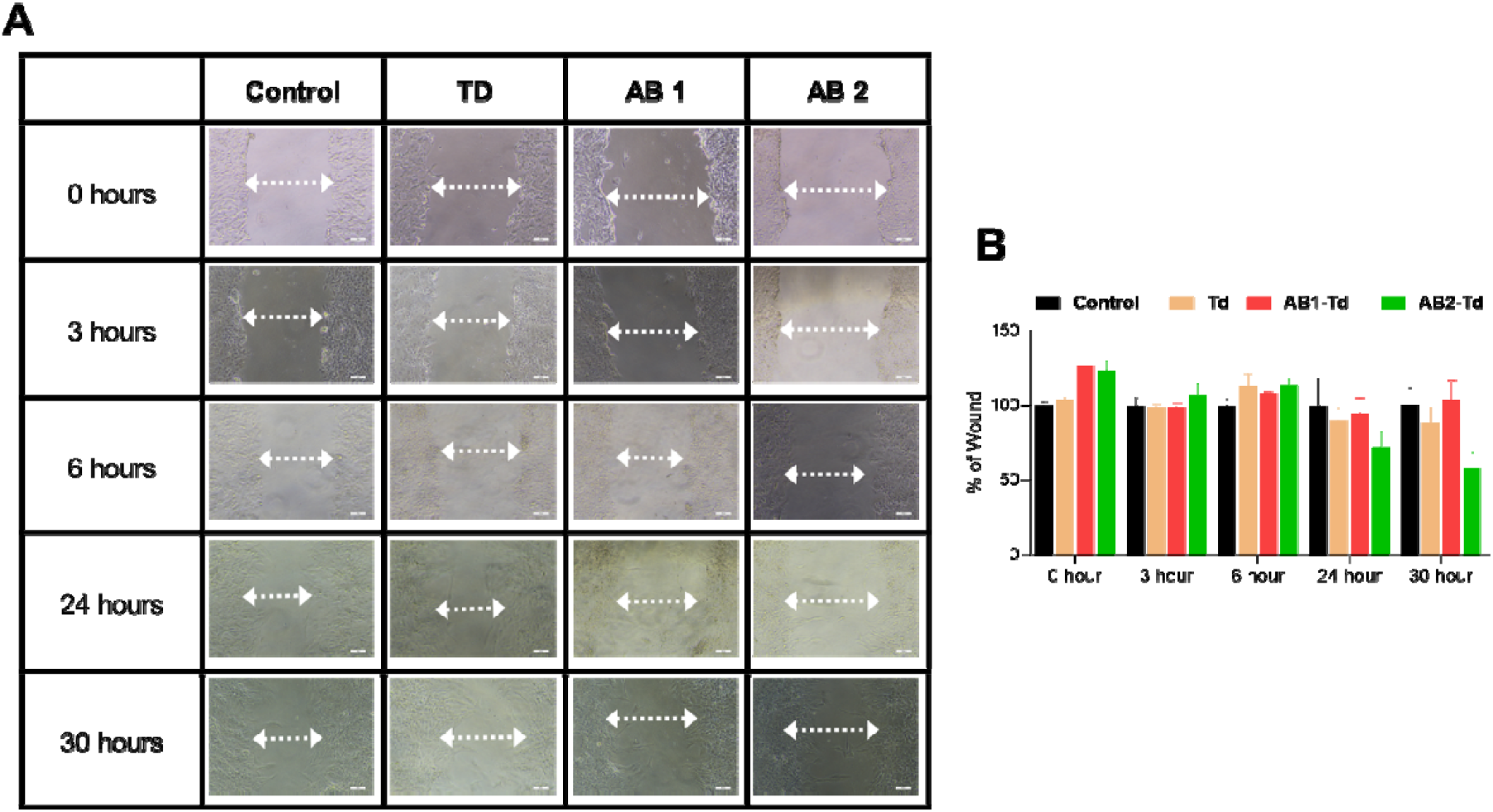
**(A)** The wound-healing process of RPE-1 cells was done at 100 nM peptide-tdDNA concentration, imaged at different time points using a Nikon phase contrast microscope The images were captured for control and cells treated with AB1/AB2-tdDNA **(B)** Quantification of wound healing in RPE-1 cells using imageJ software.

### 3.4. In vivo internalization

To evaluate the internalization capacity of our designed peptide-tdDNA system in vivo, we employed zebrafish larvae as a model system. To check the internalization, here we used Cy5-conjugated tdDNA. From **Figure 4** and quantitative analysis it is observed that our designed system has internalized effectively. This result proves, our designed short peptide-based receptor promoting peptide with tdDNA has ability to internalize in vivo.

**Figure 4.**
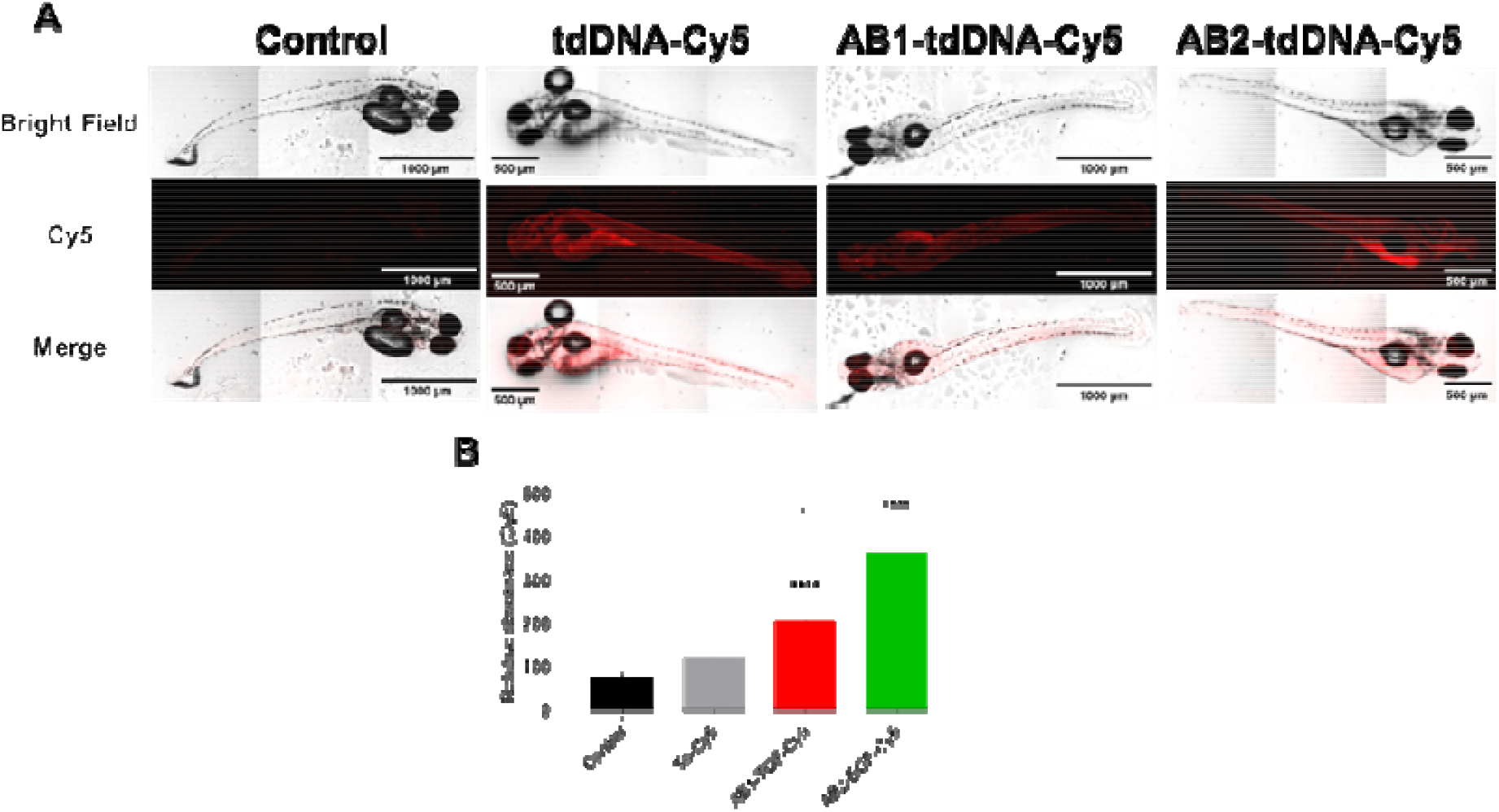
(A) Cellular internalization study of peptide tdDNAcy5 conjugates in zebrafish. (B) Quantification of cellular internalization of peptid-tdDNACy5 conjugates in zebrafish using imageJ software.

## 4. Conclusions

Though preliminary at this stage, our engineered peptide-tdDNA-based system exhibits high cellular internalization both in vitro and in vivo. Additionally, our system has EGF and TGF-β promoting short peptides, which we confirmed from a wound healing assay in RPE-1 cells. Furthermore, our designed system also confirms its successful internalization in vivo zebrafish model. Our findings on DNA-peptide based materials may broaden its potential as regenerative therapeutics. Future experiments will involve to establish specificity of uptake using pathway or receptor specific inhibitors as we as the signalling readouts using western blot analysis to prove the mechanism.

## 5. Contributions of authors

DB and AB developed the concept for the study. HNA and BP helped in optimizing experiments. DB secured the funding needed for the project. AB conducted the investigation. KK executed the in vivo model studies. AB conducted the data analysis. AK offered the necessary facilities and oversaw the in vivo experiments. SG provided resources and supervised the peptide synthesis process; VS helped in LC-MS. DB oversaw the entire project. AB wrote the initial manuscript, which was subsequently reviewed and revised by all authors.

## 6. Acknowledgment

We express our heartfelt gratitude to all the members of the DB group for their critical review of the manuscript and their invaluable insights. AB, and KK are grateful to the Department of Science and Technology, GoI for the DST Nanomission Postdoctoral fellowship. HN is grateful to the Ministry of Education, GoI for the PMRF fellowship. DB is thankful to SERB-CRG, MoES-STARS, Gujcost-DST, GSBTM, and IITGN for the research funding. A sincere acknowledgement is extended to Dr. Sharad Gupta for providing the peptide synthesis facility and to Dr. Ashutosh Kumar for the zebra fish facility. We also recognize the CIF facilities at IIT Gandhinagar for their support with Confocal and AFM facilities.

## Notes

### Competing Interest Statement

The authors have declared no competing interest.

