## Supplementary material for "Growth-factor ligand functionalization enhances cellular and *in vivo* uptake of DNA nanodevices": Supplimentary file

**Supporting Information**

(M+H)^+^

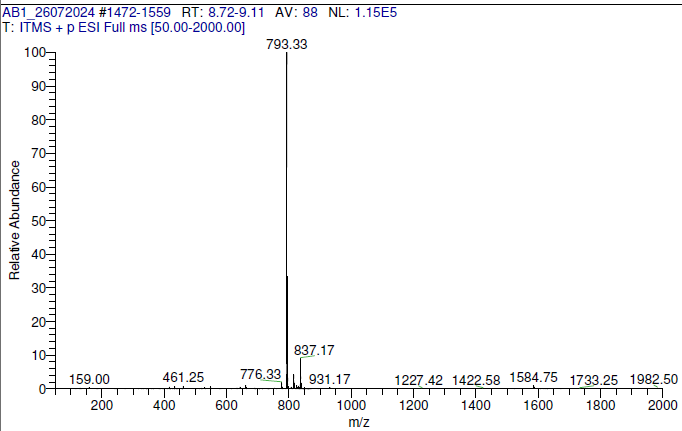

**Figure S1:** Mass spectra of AB1.

(M+3H)^3+^

(M+2H)^2+^

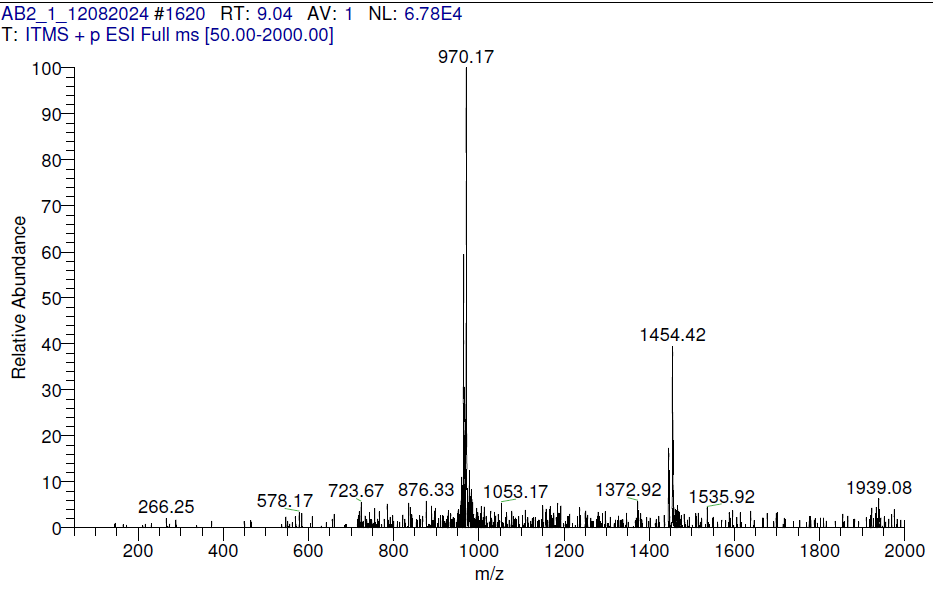
 **Figure S2:** Mass spectra of AB2.

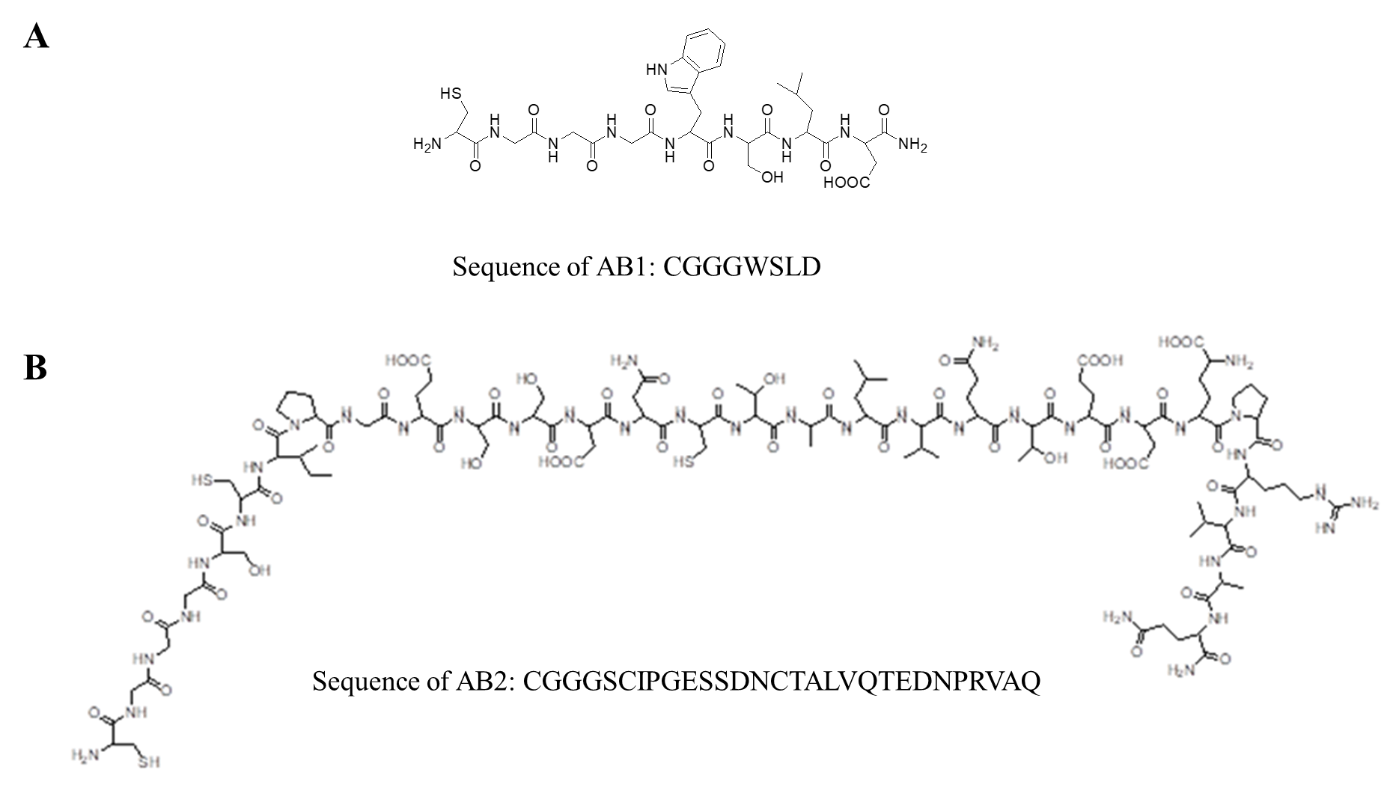

**Figure S3.** Schematics of AB1 and AB2

**Table S1:** Sequence of oligonucleotides used for the DNA tetrahedron synthesis

| **Sequence ID** | **Sequence (5’-3’)** |
| --- | --- |
| M1 - No modification | ACATTCCTAAGTCTGAAACATTACAGCTTGCTACACGAGAAGAGCCGCCATAGTA |
| M1 - Amino modification | AmC6-ACATTCCTAAGTCTGAAACATTACAGCTTGCTACACGAGAAGAGCCGCCATAGTA |
| M2 | TATCACCAGGCAGTTGACAGTGTAGCAAGCTGTAATAGATGCGAGGGTCCAATAC |
| M3 | TCAACTGCCTGGTGATAAAACGACACTACGTGGGAATCTACTATGGCGGCTCTTC |
| M4 | TTCAGACTTAGGAATGTGCTTCCCACGTAGTGTCGTTTGTATTGGACCCTCGCAT |
| M4-Cy3 | Cy3-TTCAGACTTAGGAATGTGCTTCCCACGTAGTGTCGTTTGTATTGGACCCTCGCAT |
